# Triggering mechanisms for motor actions: A mini meta-analysis and experimental data

**DOI:** 10.1101/360297

**Authors:** Li-Ann Leow, Aya Uchida, Jamie-Lee Egberts, Stephan Riek, Ottmar V. Lipp, James Tresilian, Welber Marinovic

## Abstract

Motor actions can be released much sooner than normal when the go-signal is of very high intensity (> 100dBa). Although statistical evidence from individual studies has been mixed, it has been assumed that sternocleidomastoid (SCM) muscle activity could be used to distinguish between two neural circuits involved in movement triggering. We summarized meta-analytically the available evidence for this hypothesis, comparing the difference in premotor reaction time (RT) of actions where SCM activity was elicited (SCM+ trials) by loud acoustic stimuli against trials in which it was absent (SCM- trials). We found ten studies, all reporting comparisons between SCM+ and SCM- trials. Our mini meta-analysis showed that premotor RTs are faster in SCM+ than in SCM- trials. We also present experimental data showing the effects of foreperiod predictability can induce differences in RT that would be of similar size to those attributed to the activation of different neurophysiological pathways to trigger prepared actions. We discuss plausible physiological mechanisms that would explain differences in premotor RTs between SCM+ and SCM-trials.

## Introduction

An unexpected startling acoustic stimulus (SAS) delivered during the preparation for a motor action can trigger the prepared response at a latency that is much shorter than normal, a phenomenon termed the StartReact effect [1]. Because the StartReact effect can reduce deficits in motor initiation and execution in some neurological conditions [2-7], it has the potential to be applied in movement rehabilitation interventions. However, there is ongoing debate around what exactly constitutes a true startle response and how the StartReact effect can be differentiated from other well-known phenomena such as the stimulus intensity effect [8]. This debate is theoretically relevant because it concerns our basic understanding of how motor actions are prepared and initiated by the central nervous system (CNS).

Carlsen and colleagues proposed that to ascertain that the neural mechanisms responsible for the StartReact effect are activated, it is essential to observe sternocleidosmastoid (SCM) activity [9, 10]. Thus, trials where responses are triggered by unexpected startling acoustic stimuli must be divided into trials in which the SAS elicited a SCM response (SCM+ trials) and trials where it did not (SCM–). Although we have highlighted the difficulties with interpretation of data based on a strict adherence to this criterion elsewhere ([11], it has been generally accepted that SCM activity is important to study the StartReact effect. It has even been proposed that, when analyzing the StartReact effect, trials should be discarded when no activity is detected in this muscle, as the physiological mechanism for movement triggering would be different [12]. However, not all studies separating trials based on SCM activity (SCM+ and SCM-) report statistically reliable differences in reaction times (RT) [5, 13, 14] and in some instances the StartReact effect (very short RTs) appears to be observed even with no SCM activity at all [15], bringing into question the hypothesis that there is a special startle-evoked mechanism at play only in SCM+ trials. In general, however, it has been proposed that SCM+ trials comprise a distribution of RTs that are on average shorter than those observed for SCM- trials. Alternatively, Marinovic and colleagues have proposed that differences in SCM+ and SCM- activity may be explained by random fluctuations in preparatory activity from trial to trial [11, 16], which could lead to both a reduction in RT and a higher probability of detecting SCM activity when preparation levels are high.

To the best of our knowledge, no study has systematically analyzed the differences in RT between SCM+ and SCM- trials in StartReact studies. Here, we conducted a mini meta-analysis to: **a)** verify whether SCM activity indeed indicates differences in the speed of triggering the motor plan and, if so, **b)** estimate the magnitude of this difference so that we design studies with appropriate sample sizes. Additionally, we sought to determine whether factors contributing to fluctuations in motor preparation levels such as foreperiod variability (see [17]) could be associated with differences in the results of published studies. The results of our meta-analysis showed that SCM activity is correlated with faster reaction times. The meta-analysis also demonstrates that foreperiod variability makes an important contribution to the observation of the StartReact effect. To further examine the impact of foreperiod variability on reaction times elicited by SAS, we conducted an experiment varying the temporal certainty of the imperative stimulus. The results showed that temporal certainty does impact the reduction of reaction times to SAS, which could explain differences between SCM+ and SCM- trials.

## Mini meta-analysis

### Methods

We conducted a Scopus search of research articles citing either Valls-Solé et al.’s [1] seminal paper on the StartReact effect or Carlsen et al.’s paper [9] suggesting that responses for which SCM activity was observed were triggered differently from responses without such activity. Because we wanted to more precisely estimate the magnitude of the “true” StartReact effect in comparison to stimulus intensities effects, our analysis only included experimental studies that reported premotor reaction times in SCM+, SCM-, and control trials (trials for which the SAS was replaced by either a less intense acoustic or visual go signal). We performed separate meta-analyses comparing the reaction time means of both SCM+ vs. SCM- trials and SCM- vs. control trials. Meta-analyses were performed using the random effects model, calculating Q-statistics as an indicator of heterogeneity. Standardized effect sizes for the mean change in premotor reaction time were calculated using raw score standardization with heteroscedastic population variances proposed by Bonett [18]. For all meta-analyses we used the R package METAFOR [19]. Sample size calculations to obtain a power of 80% and an a-error of 0.05 were performed using G*Power 3.1 [20].

### Results

The initial search yielded 210 research articles, of which ten were included for the meta-analysis. Manuscripts were excluded from analysis if (1) SCM activity was not recorded or (2) a comparison between trials in which SCM+ and SCM- was not conducted (e.g. only SCM+ trials were analyzed). For one of the included studies, additional data was kindly provided by the authors [21] to allow the computation of correlation coefficients between change scores. Means and standard deviations of premotor reaction times were provided in the text of the papers or extracted from plots using WebPlotDigitiser 3.8. For six studies, we could not obtain the data to calculate correlation coefficients between change scores [3, 9, 10, 22-24]. Sensitivity analysis using correlation coefficients from 0.05 to 0.95 showed only small differences in our estimates (≈ 2 ms), thus, we adopted a correlation coefficient of 0.5 for the studies for which data was not available.

#### Differences between SCM + and SCM - trials

The difference in premotor reaction time between SCM+ and SCM- trials was −15.5 ms (95% CI [-21.81, −9.3], t = 5.62, p = 0.0003) and the heterogeneity was also statistically significant (Q test χ2 = 21.64, df = 9, p = 0.01). As shown in Figure 1, the direction of the effect is consistent across all studies; that is, responses are typically released earlier when SCM activity is detected. This is consistent with the proposal that truly startled responses (SCM+) should be faster than non-startled responses (SCM-). An estimate of the standardized effect size [18] suggests that this effect is large to medium (point estimate: −0.79, 95% CI [-1.14,-0.44]). A sample size estimate based on the average effect size (i.e., one-tailed t-test) indicates this effect could be detected with 12 participants. However, a more conservative estimate of the required sample size to detect a medium effect size (i.e., close to the lower bound of the confidence interval of the standardized effect size) would be 34 participants, indicating most studies to date would be underpowered to detect medium sized effects. The estimated average premotor reaction time in SCM+ trials based on the nine studies using simple RT tasks was 89.8 ms (95% CI [82.9, 96.7]).

**Figure 1:**
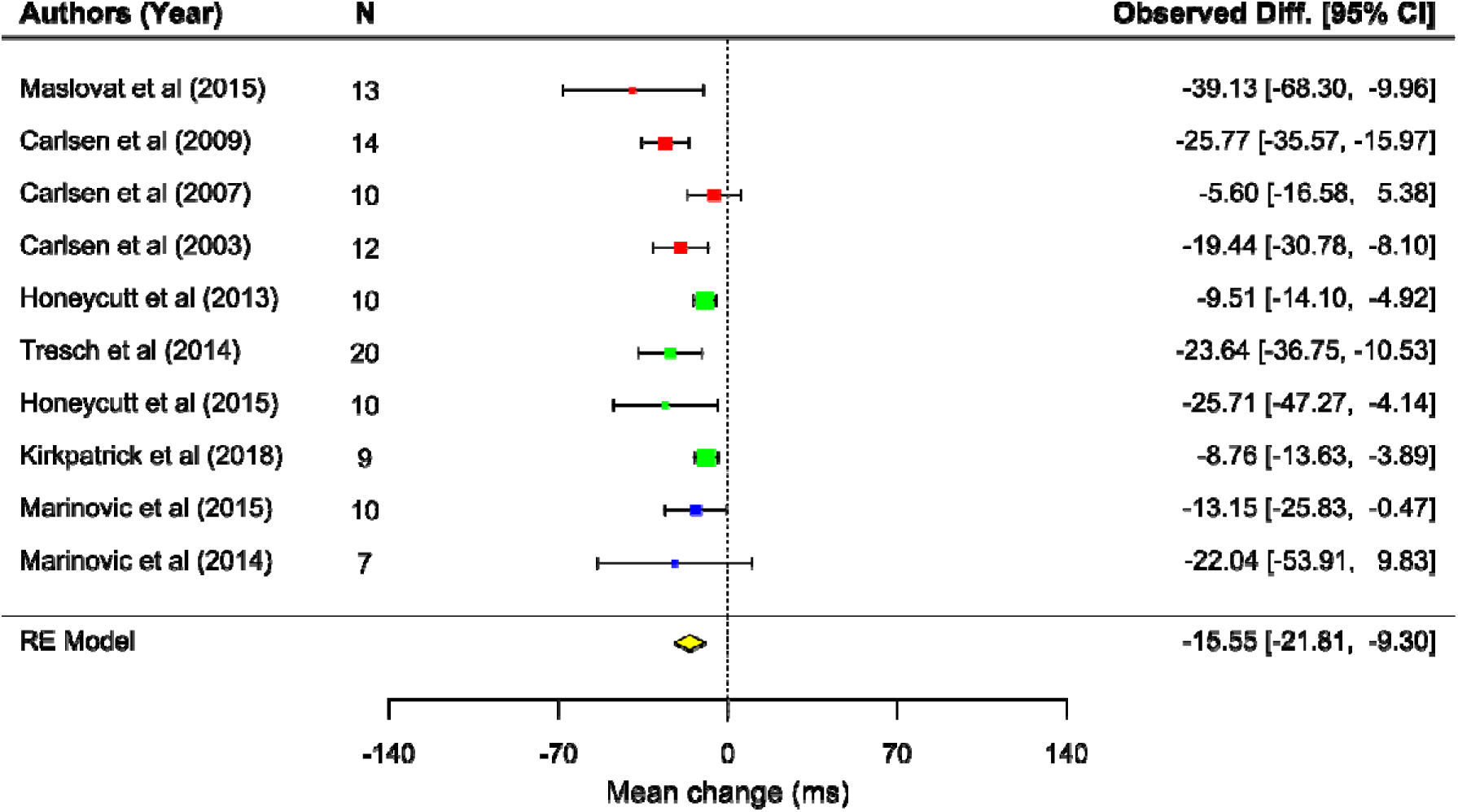
Forest plots and meta-analysis of studies comparing SCM+ and SCM- trials (n = 115).

#### Differences between SCM- and Control trials

Because three experiments used either a visual go-signal in control trials [10, 13] or no go-signal [14], we limited our comparisons between SCM- and control trials to the seven experiments which used a soft go-signal (≈ 80dBa) in control trials. This analysis showed that the difference in premotor reaction time between SCM- and control trials across these studies was −45.59 ms (95% CI [-62.60, −28.58], t = 6.95, p = 0.0006) but the heterogeneity was also statistically significant (Q test χ2 = 28.19, df = 6, p < 0.0001, see Fig. 2). The standardized effect size [18] indicates that this effect is large (point estimate: −1.63, 95% CI [-2.18,-1.08], Q test χ2 = 9.85, df = 6, p = 0.13), and can be detected - one-tailed t-test - with small sample sizes (≈ 7 participants; based on the lower bound of the effect size confidence interval). As it can be seen in Table 1, however, the raw mean differences in premotor reaction times of the studies can be divided into two subgroups according to the variability of the foreperiod (time between warning-signal and the go-Signal). For the three studies with lower foreperiod (≤ 500 ms) variability (see Table 1 and Figure 2), the difference in the mean premotor RT between SCM- and control trials was −32 ms (95% CI [-41. 5, −22.5], t = 14.45, p = 0.004; Heterogeneity: Q test χ2 = 0.94, df = 2, p = 0.62). The standardized effect size showed that this effect is medium to large (point estimate:-1.45, 95% CI [-2.28, −0.62], Q test χ2 = 1.41, df = 2, p = 0.49). In studies with relatively larger foreperiod (≥1000 ms) variability, the difference between SCM- and control trials was - 58.5 ms (95% CI [-86.03, −30.99], t = −6.76, p = 0.006; Heterogeneity: Q test χ2 = 9.84, df = 3, p = 0.019), close to double the estimate obtained for the lower variability subgroup as shown in Figure 2. Here the standardized effect size [18] was the largest with a point estimate of −1.99, but with a statistically significant heterogeneity (Q test χ2 = 4.25, df = 3, p = 0.023), explaining the large confidence interval for this effect size (95% CI [-3.48, 0.50]). It is worth mentioning that the difference between Higher and Lower foreperiod variability subgroups in terms of RT latencies in controls trials is diminished by the lower RTs in Kirkpatrick et al.’s [25] study where participants practiced for 10 days, and we analyzed here only data from the last day when RTs were reliably faster than on the 1st day of the study. If we exclude Kirkpatrick et al.’s data from our analysis of subgroups, the difference between Higher and Lower foreperiod variability subgroups is more than doubled (Higher = 68.1 ms, Lower = 32 ms).

**Figure 2:**
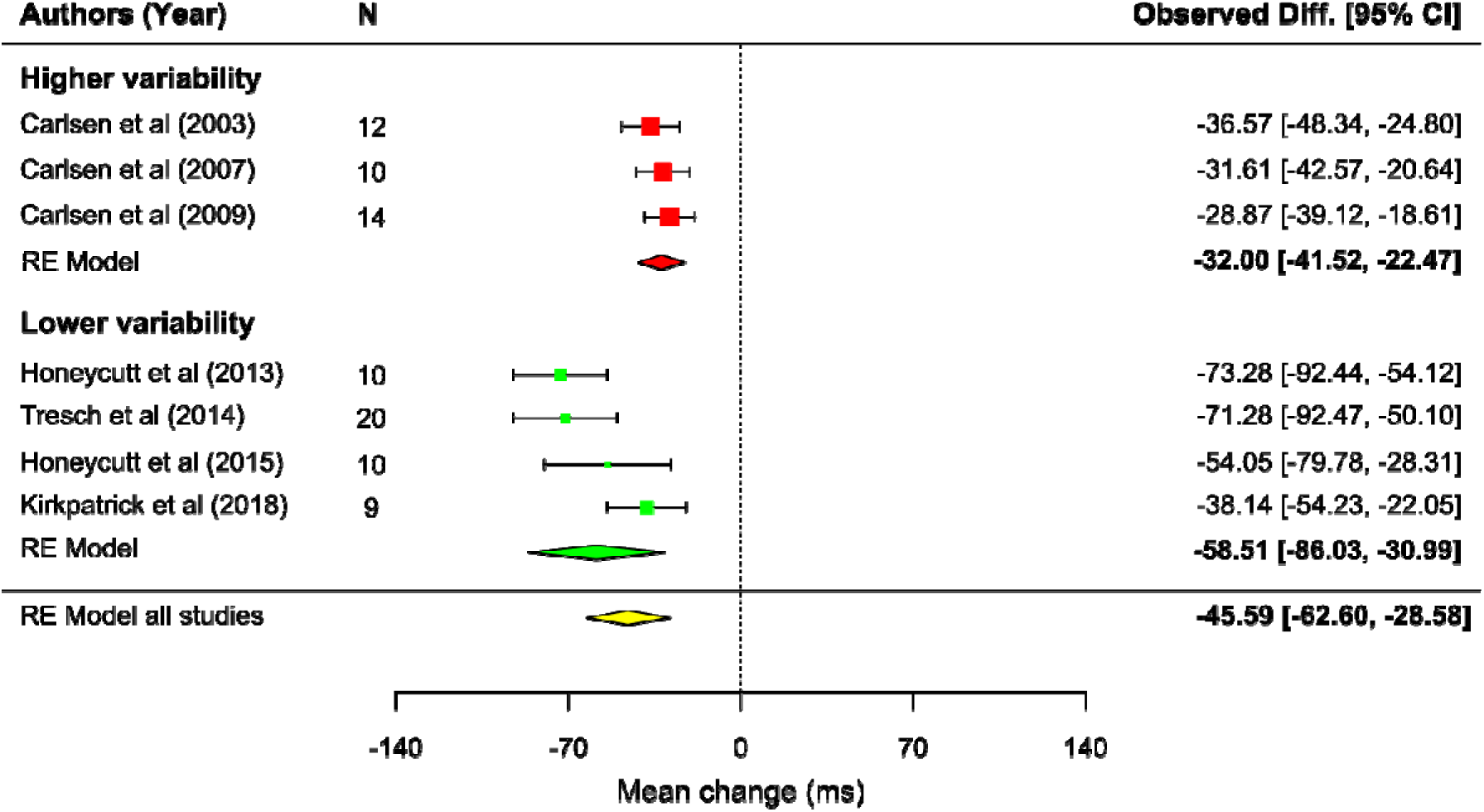
Forest plots and meta-analysis of studies comparing SCM- and control trials in which the go-signal was acoustic (n = 85).

**Figure 3:**
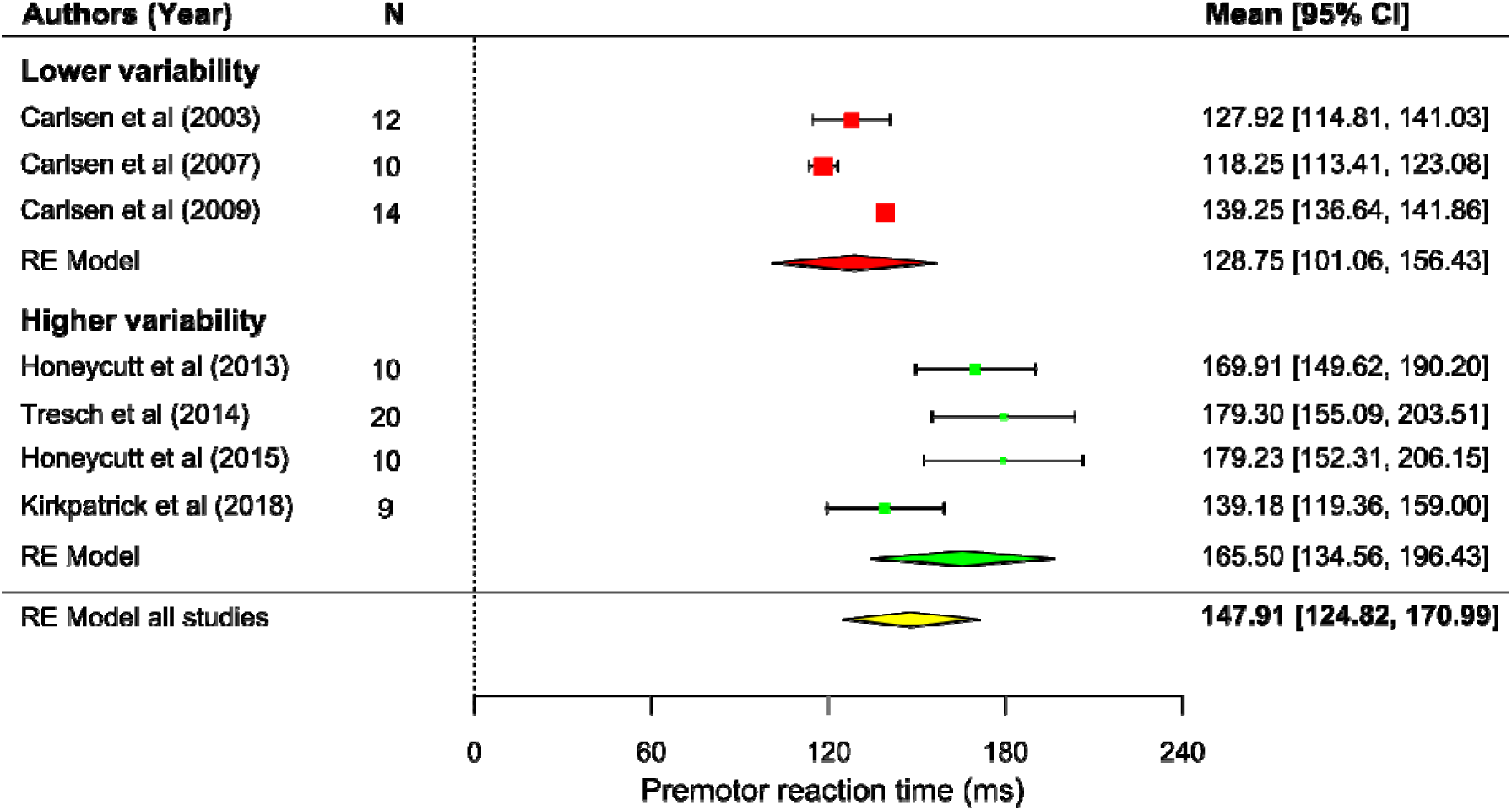
Forest plots and meta-analysis of studies showing the average premotor reaction in control trials (n = 85).

**Figure 4:**
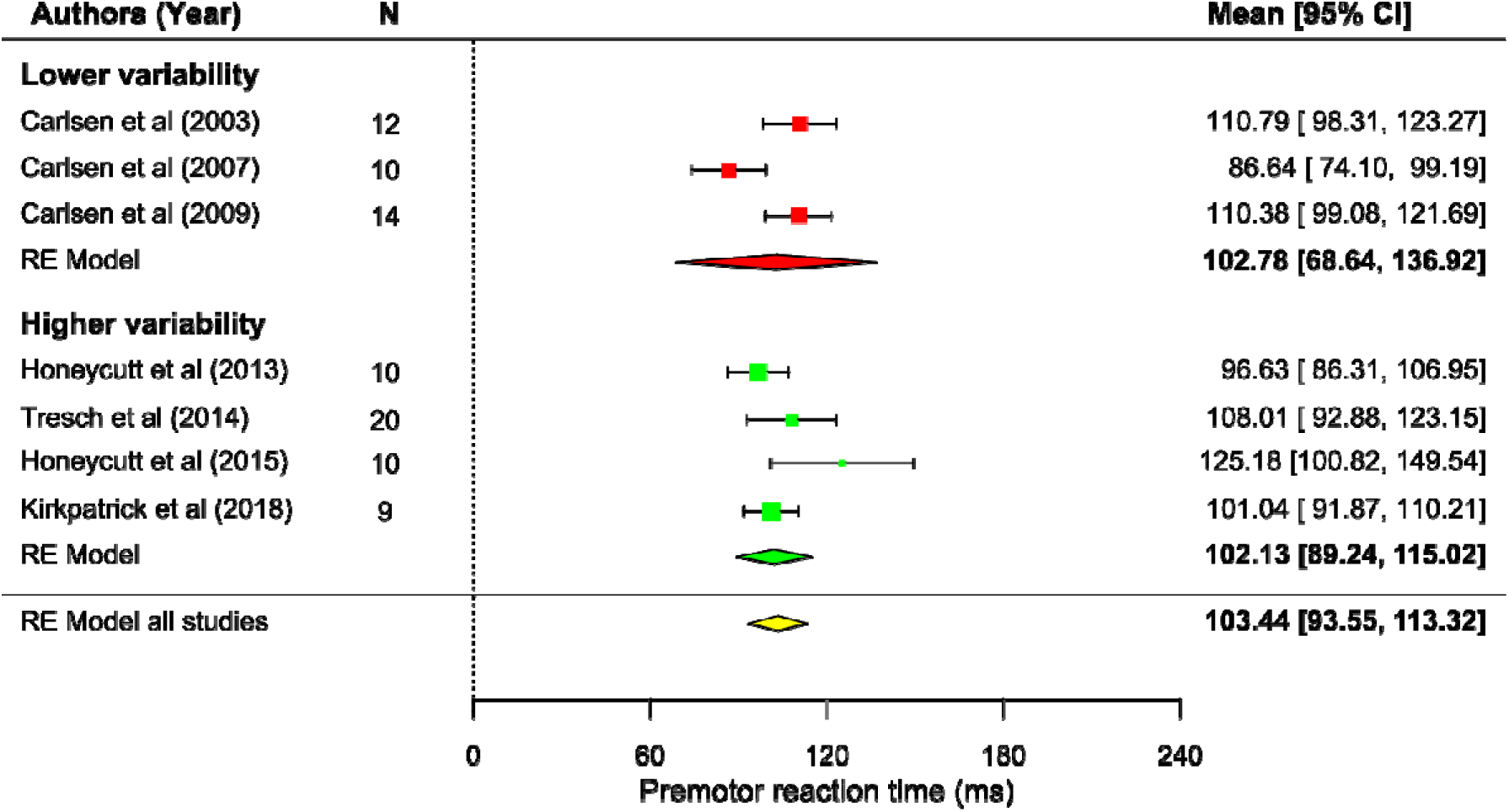
Forest plots and meta-analysis of studies showing the average premotor reaction in SCM- trials (n = 85).

**Table 1:** Average time between soft warning-signal and loud go-signal (foreperiod) and its variability.

| Authors (Year) | Average foreperiod (sec.) | Foreperiod variability (+/-seconds) |
| --- | --- | --- |
| Carlsen et al. (2003) | 2.5 | 0 |
| Carlsen et al. (2007) | 2.5 | 0 |
| Carlsen et al. (2009) | 2.5 | 0.5 |
| Honeycutt et al (2015) | 2.5 | 1 |
| Tresch et al (2014) | 2.5 | 1 |
| Honeycutt et al (2013) | 2.5 | 1 |
| Kirkpatrick et al (2018) | 2.5 | 1 |

#### Experiment

Our meta-analysis indicated that responses with SCM activity are initiated significantly earlier than responses without it, and also revealed that foreperiod variability has a substantial effect on the magnitude of the difference between responses elicited in control and probe trials (SAS presented). It is well known that foreperiod manipulations can strongly affect the level of preparation for an action and, consequently, modulate RT [17]. However, Niemi [26] showed that foreperiod effects are reduced when the IS is an intense signal in the auditory modality, suggestive of a facilitatory effect of phasic arousal over the neural activity of circuits responsible for the initiation and execution of the voluntary response [17, 27]. This reduction of foreperiod effects when using an intense acoustic go-signal may reflect the activation of a specific physiological mechanism thought to be involved in the StartReact effect [28]. In what follows, we sought to revisit this issue and determine whether systematic fluctuations in the level of preparation for action - as indexed by RT measurements - can be observed when the IS is a strong signal in the auditory modality (SAS). This allowed us to evaluate our hypothesis that random fluctuations in preparatory activity from trial to trial [11, 16] are associated with reductions in RT in StartReact studies.

## Methods

### Participants

Twenty-four volunteers (mean age 19.5 years old, SD = 3.17; 14 female) participated in the experiment. All of them stated that they were right-handed and had normal of corrected to normal vision. Participants gave informed consent prior to commencement of the study, which was in accordance with the Declaration of Helsinki and approved by the local Ethics Committee of the University of Queensland.

## Task

To induce systematic variations in preparatory states over time during a trial, we used a marked reaction time task (see [29]) where the IS (a soft or a SAS) was presented together with one of three visual cues (visual time markers). More precisely, we presented a sequence of four brief flashes (50 ms, red square, 200 x 200 pixels) displayed 600 ms apart as shown in Figure 5. The first flash served as the warning signal (WS), whereas subsequent flashes marked the potential temporal location of the IS (600, 1200, or 1800 ms after the WS), and eliminated uncertainty about the temporal aspect of the task. With this task, we expected that RT would decrease as a function of the evolving conditional probability of the presentation of the IS during a trial when its temporal location was unpredictable [17, 29]. In other words, reductions in RT were expected to occur as a function of the increasing probability of the IS over time. Participants performed this task in two blocks: a) predictable, and b) unpredictable. In the predictable block, participants were informed that the IS (soft or SAS) would be presented together with the 2^nd^, 3^rd^, or 4^th^ flash before the trials began. The order of marker presentation was randomized across participants, and all trials for each IS-marker pairing (2^nd^, 3^rd^ or 4^th^) were presented sequentially. Note that to avoid false starts, we also presented catch trials in which the IS was not presented. In the unpredictable block, participants were informed that the IS could be presented together with any of the three temporal markers or not presented at all (catch trials). In each block of trials, participants performed 27 trials (9 trial/marker) in which the SAS was presented, 27 catch trials (9 trials/marker), and 108 control trials (36 trials/marker). The order of the blocks was counterbalanced across participants to avoid sequencing effects.

**Figure 5:**
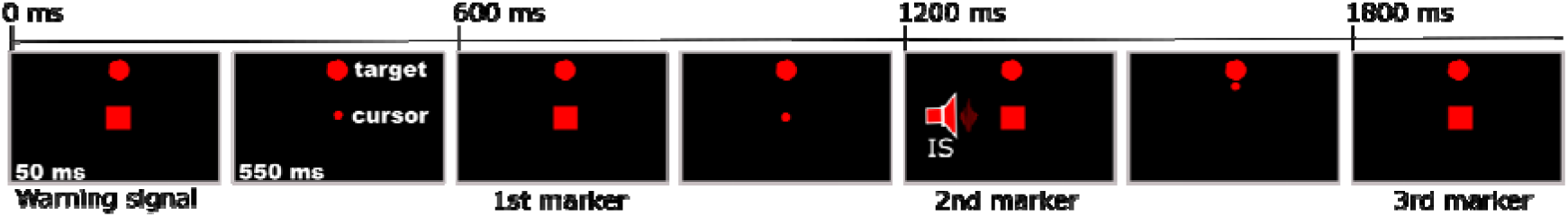
Sequence of events during a trial. The first flash served as a warning signal (WS) and lasted for 50 ms. Subsequent flashes were presented 600 ms apart. The second, third, and fourth flashes also lasted 50 ms, and represented the 1st, 2nd, and 3rd temporal markers. The target was presented simultaneously with the WS and served to motivate participants to produce at least 20N of force in all trials. A moving cursor controlled by the participants represented the forces applied during the trial. To produce the minimum force required participants had to make the cursor intersect the target. Participants were asked to make an abduction of the wrist (radial deviation) as fast as possible upon hearing the imperative stimulus (IS). In the illustrative figure, the IS was presented in synchrony with the 2nd marker, and upon response onset the cursor moved towards the target.

## Procedures and Design

Participants sat in a chair in front of a 22-in Samsung LCD monitor (60 Hz refresh rate, 1280 x 1024 resolution) approximately 1 m in front of them. The task required participants to make isometric abductions of the wrist (radial deviation) as fast as possible upon hearing the IS (see Figure 5). Participants had their right hands snugly fit into a custom-built device (see [30, 31]) that held the hand and forearm in a neutral position throughout the experiment. To standardize the level of force produced in each trial, participants were asked to move a circular cursor from the centre of the monitor to a target presented at 90° in relation to the cursor origin (see Figure 5). To move the cursor to the target, participants had to apply a contraction equal to 20Ns with their wrists. Forces were measured by a six-degree of freedom force/torque sensor (JR3 45E15A-I63-A 400N60S, Woodland, CA). Veridical feedback on RT was provided on the monitor screen after all control trials to encourage fast responses and also avoid anticipatory reactions. Thus, any response with a RT of more than 200 ms was followed by the text “Too slow”, whereas any response with a RT of less than 50 ms was followed by the text “Too quick”. Responses that fell within these intervals were followed by a “Good timing” message. When the IS was intense (a SAS), the message displayed was always “Good timing” irrespective of the actual RT value. Catch trials were followed by the message “No movement required”.

Prior to the experimental trials in each bock, participants performed 15 practice trials to familiarize themselves with the task. Acoustic stimulation was presented three times during familiarization. Visual stimuli were generated with Cogent 2000 Graphics running in MATLAB 7.5.

## Auditory stimuli

The auditory stimuli were generated with MATLAB and presented binaurally through high fidelity stereophonic headphones (Seinheiser model HD25-1 II; frequency response 16Hz to 22kHz; Sennheiser Electronics GmbH & Co. KG, Wedemark, Germany). The input signal to the headphones had a bandwidth of approximately 10 Hz–30 kHz. The soft auditory IS was a 50 ms pure tone (500 Hz) with a peak loudness of 65 dBa, whereas the SAS was a broadband white-noise (rise/fall time shorter than 1 ms) with a peak loudness of 114 dBa. Sound intensity was measured with a Bruel and Kjaer sound level meter (type 2205, A weighted; Brüel & Kjaer Sound & Vibration Measurement, Naerum, Denmark) placed 2 cm from the headphone speaker.

## Data analysis

The main variable of interest was reaction time (RT). RT was defined as the difference between the time of movement onset and the time the IS was presented. To calculate movement onset times, we first transformed the two-dimensional screen coordinates and filtered it using a low-pass second order Butterworth filter with a cut-off frequency of 10 Hz. Movement onsets were estimated from the tangential speed time series (derived by numerical differentiation of the filtered cursor position data) using an algorithm proposed by Teasdale et al. [32]. Although it is typical to report pre-motor reaction times in the StartReact literature (see our mini meta-analysis above), our RTs measurements using the torque data add only a small delay to our estimates of movement onset time. More precisely, based on data from Marinovic et al. [33] (26 participants in Experiment 2), the average introduced delay is about 19.3 ms (SD = 4.9), allowing us to estimate how much quicker pre-motor RTs should be based on our RT data. We also analyzed the percentage of false starts as a function of IS timing and predictability. A response was considered a false start if the participant’s rate of force development - the first derivative of the forces applied on the torque sensor - surpassed 10% of the median rate of force development observed in trials with a soft IS. Trials in which RT was lower than 50 ms (anticipatory reactions) or larger than 1000 ms (inattention to the IS) were discarded. Across all participants, approximately 2.9% (SD = 5) of all trials were excluded from further analysis based on this criterion, and the inclusion of all trials did not change the qualitative pattern of results.

The statistical data analysis was conducted in R ([34]) using the lmer function from the lmerTest package [35]. The analysis was separated into two phases. First we analyzed the median RT using a 2 (Predictability: Predicable vs. Unpredictable) x 2 (IS intensity: soft IS vs. SAS) x 3 (IS time: 1st, 2nd, and 3rd marker) linear mixed model. For this analysis, we employed the Sattethrwaite approximation [36] to calculate F-tests and estimate p-values for the main effects and their interactions. Predictability, IS intensity and IS time were treated as fixed factors, whereas participants were treated as a random factor into the model. The percentage of false starts was analyzed using a permutational analysis of variance using the *ezPerm* function (*ez* Package), and had Predictability and IS time as factors. In the second phase of our analysis, we fitted cumulative distribution functions (CDF) to the data of all participants in both blocks of trials when the IS was intense (SAS). Next, we recorded the 30th and 60th percentiles of each CDF for each participant. These values were then entered into a 2 (Predictability: Predicable vs. Unpredictable) x 2 (Percentile: 30th vs. 60th) x 3 (IS time: 1st, 2nd, and 3rd marker) linear mixed model. Predictability, RT Percentile, and IS time were treated as fixed factors, whereas participants were treated as a random factor into the model. The rationale for this analysis relies on the assumption that the distribution of RTs in response to SAS is bimodal (see [11]), Figure 4), reflecting the activation of different neurophysiological pathways to trigger prepared responses. Note that, typically, the method of choice to determine if trials were triggered via the mechanism responsible for the StartReact effect is the presence of SCM activity [21]. However, given that responses can be elicited rather quickly by a SAS in the absence of SCM activity [15], and slow responses can be observed when SCM activity is detected [11], one can conclude that SCM activity is neither necessary nor sufficient to determine whether the StartReact effect was observed. Therefore, separating trials by their latencies is likely to be more indicative of a mechanism that bypasses or activates specific mechanisms in the central nervous system than relying on surface EMG (see also [37]). To decide which percentiles would be representative of values obtained when using SCM to separate trials (SCM+ and SCM-), we fitted a CDF to data kindly provided by Honeycutt et al. (2013) (grasp task included in the meta-analysis) where a statistically reliable effect was observed. For SCM+ trials, Honeycutt and colleagues reported a mean latency of 87 ms, which matched closely (86 ms) the 30^th^ percentile estimate using a CDF. For SCM- trials, they reported a mean latency of 96 ms, which approximately matched (94 ms) the 60^th^ percentile using a CDF. To further examine the impact of presenting the intense IS (SAS) while preparation levels were expected to vary over time, we also estimate the slope of linear regressions using a bootstrapped procedure (5000 iterations). If the earliest responses (30th percentile) were triggered via a distinct mechanism responsible for the StartReact effect, the confidence interval of the bootstrapped distribution should include zero. In contrast, if preparation levels could modulate the latency of the fastest responses across the three IS times, the confidence interval of the bootstrapped slope distribution should not include zero.

Effect sizes are given as likelihood ratios (LR) and were calculated contrasting the null or main effect models against relevant models of interest (single main effects or interactions). All experimental data can be accessed via this link: https://data.mendeley.com/datasets/m7j6zkfhz8/draft?a=d59740b0-b029-4c62-8ee4-3d9b70c79d15.

## Results

### Initial analysis

As expected, there was a main effect of IS intensity on RT (F_(1,253)_ = 130.08, p < .0001, LR = 63.4), indicating that responses were faster when the IS was intense (SAS). There was also a main effect of temporal predictability of the IS (F_(1,253)_ = 106.49, p < .0001, LR = 50.7), demonstrating that RTs were shorter when participants knew the likely time of appearance of the IS. The main effect of IS timing was also statistically reliable (F_(2,253)_ = 39.61, p < .0001, LR = 36.7), indicating responses tended to become faster as the IS appeared later in the trial. As shown in Figure 6A, this main effect was qualified by a significant interaction between IS predictability and IS timing (F_(2,253)_ = 18.66, p < .0001, LR = 24.36), suggesting the effect of IS timing was stronger in the unpredictable block of trials.

**Figure 6:**
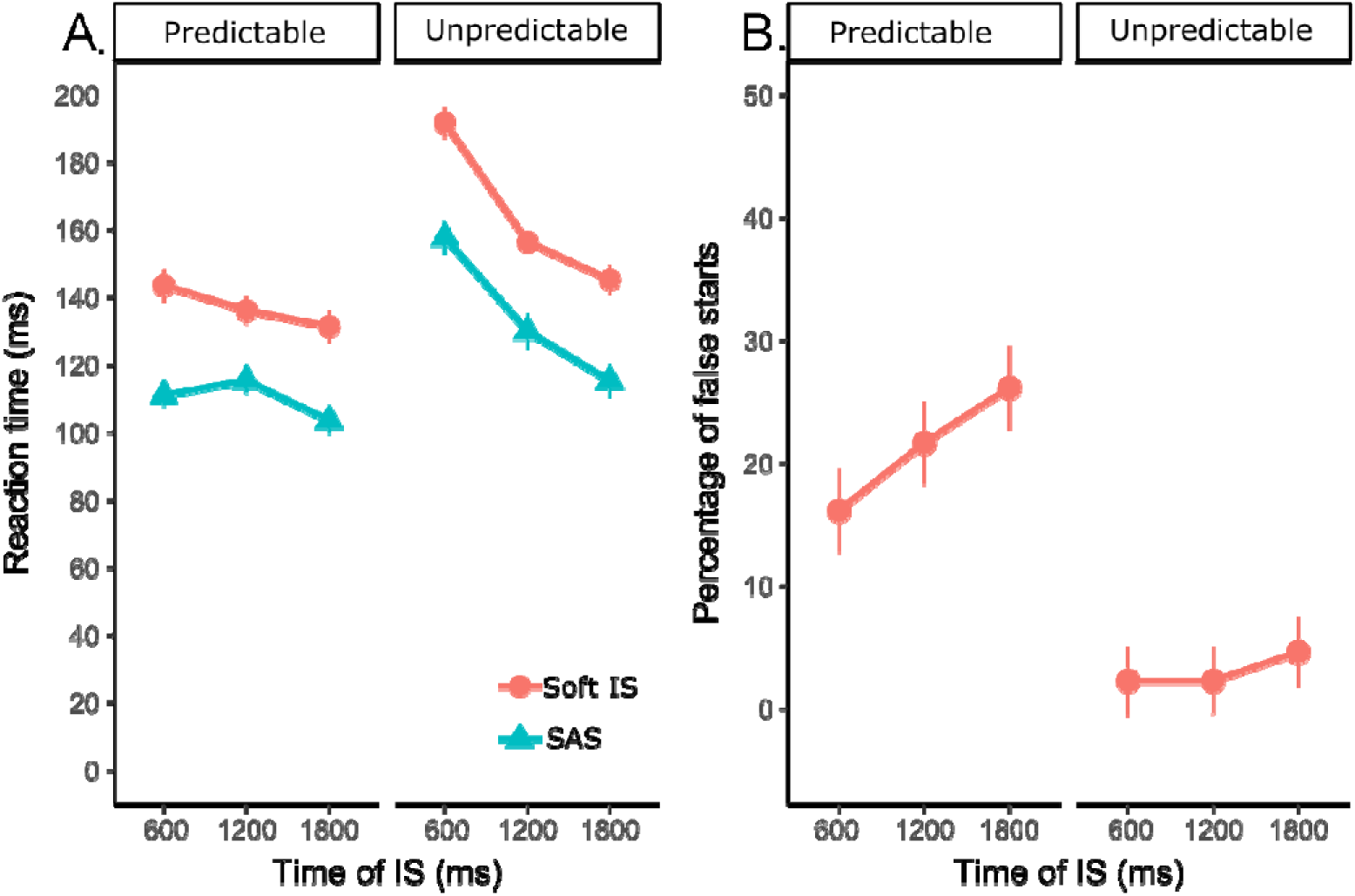
**A.** Reaction time. **B.** False starts. Error bars represent the within participants standard error of the mean [38].

There was a statistically reliable increase in the percentage of false starts as a function of IS time in both blocks of trials (main effect of IS timing: p = 0.032). This analysis also indicated a main effect of IS predictability (p < .001), indicating that participants had more false starts (≈20%) when the temporal predictability of the IS was high as shown in Figure 6B. The apparent interaction between IS time and predictability failed to reach statistical significance (p=0.094).

### Fast vs. Slow responses to SAS

As shown in Figure 7, the estimated RT at the 30^th^ percentile of the CDF were faster than those at the 60^th^ percentile (main effect of RT percentile: (F_(1,257)_ = 11.94.3, p < .0001, LR = 35.2). The pattern of results is similar to that observed when contrasting the soft and intense IS (see Figure 6A). More specifically, we observed a statistically reliable main effect of IS predictability (F_(1,257)_ = 75.1, p < .0001, LR = 54.5), and also a main effect of IS time (F_(1,257)_ = 70.36, p < .0001, LR = 38.5). The interaction between IS predictability and IS time was also statistically significant (F_(1,257)_ = 28.6, p < .0001, LR = 22.4), again suggesting that the effect of IS time was more pronounced in the unpredictable than the predictable block of trials.

**Figure 7:**
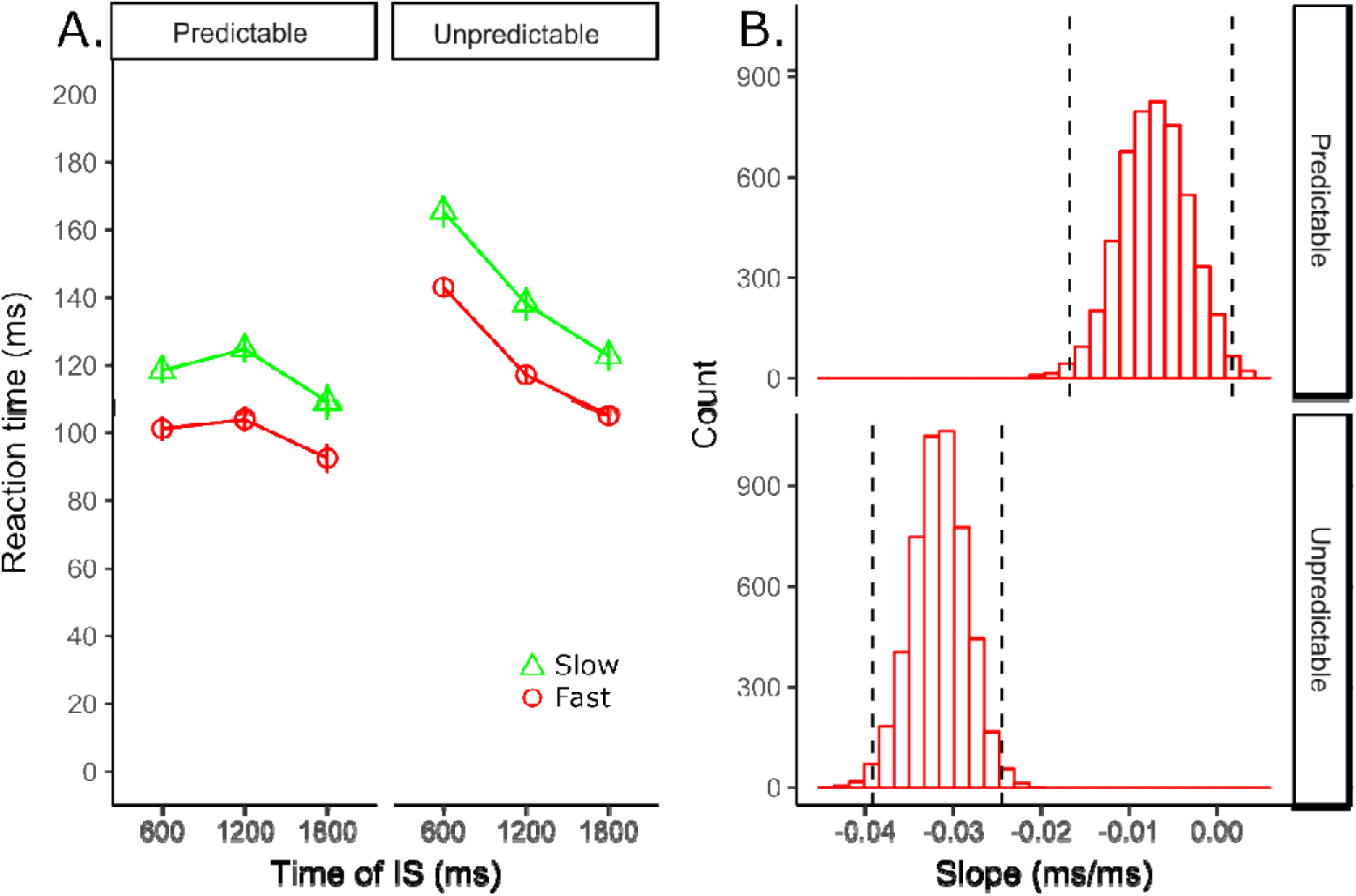
**A:** Reaction time for fast (30^th^ percentile) and slow (60^th^ percentile) as a function of IS time in both blocks of trials (Predictable and Unpredictable) for responses elicited by the intense IS (SAS = 114dBa). Error bars represent the within-participants standard error of the mean [38]. **B:** Histograms of the bootstrapped distributions of the slopes for fast RTs. Dashed black lines indicate the boundaries of the 98% confidence intervals of the linear regression slope.

To further examine whether changes in preparation levels over the course of a trial could impact the RT to an intense IS, we bootstrapped the slope of a linear regression for the RT estimates in the 30^th^ percentile for both blocks of trials. As shown in Figure 7B, the mean slope of the bootstrapped linear regression in the unpredictable block was much further away from zero (mean = −0.031, 98%CI [-0.039, −0.024]) than that obtained for the predictable block of trials (mean = −0.007, 98%CI [-0.017, 0.001]), suggesting IS predictability can also affect the latency of responses initiated within very short latencies. Note that the estimates for the unpredictable block of trials changed little when we calculated the slopes using lower percentiles (20^th^: mean = −0.026, 98%CI [-0.032, −0.020]; 10^th^: mean = −0.022, 98%CI [-0.027, −0.016]).

## General discussion

### Meta-analysis

Our mini meta-analysis revealed that premotor RTs are indeed shorter in SCM+ than in SCM- trials. The estimated magnitude of this effect across the studies is on average large enough to entertain the possibility that motor programs are triggered via a pathway that bypasses some cortical areas of the brain [1, 24], but not large enough to completely rule out cortical involvement [11]. Although the meta-analysis showed that the effect associated with SCM activity is robust across studies, the heterogeneity test approached significance and depended on the correlation coefficient chosen for the studies for which data were not available: an increase in the correlation coefficient from 0.5 to 0.56 resulted in a statistically significant heterogeneity test. This possible heterogeneity could reflect differences in the magnitude of the loud acoustic stimulus, delivery mode (headphones *vs.* speakers), and/or the level of preparation for action as a function of the uncertainty of the timing of the go-signal. In this regard, it is clear from our meta-analysis comparing SCM- and control trials that foreperiod variability is critical for the average difference among the studies analyzed. As shown in Figure 2, the differences between SCM- and control trials can be separated into 2 subgroups according to foreperiod variability. For the studies with lower foreperiod variability (see Table 1), the average premotor RT in control trials (soft go-signal) was 128.7 ms (see Figure 3). In contrast, the average premotor RT in control trials in studies with relatively larger foreperiod variability was 165.5 ms (see Figure 3). These results suggest that one cannot distinguish among intersensory facilitation [39], stimulus intensity [8, 40], and StartReact [1] effects based on the difference between premotor RTs in control trials (soft-go signal, ≈ 80dBa) and probe trials (Loud go-signal), as an increase in foreperiod variability can substantially increase the premotor RT in control trials (≈37 ms change in RT across the studies analysed here). This is consistent with previous reports on the effects of foreperiod variability in auditory RT tasks (e.g. [17, 41]). In contrast, although premotor RTs in response to loud acoustic stimuli (>114 dBa) are somewhat heterogeneous, differences in premotor RT as a function of foreperiod variability are clearly reduced (i.e., differences in RT are not as clear for low and high variability foreperiods), showing similar average estimates and overlapping confidence intervals for the subgroups (see Figure 4).

#### Why would responses be initiated earlier when SCM activity is detected?

Maslovat et al. [10] proposed that the detection of SCM activity indicates that a more direct neural circuit, associated with the startle reflex, was responsible for involuntarily triggering the prepared response. Thus, when responses occur without SCM activity (SCM-), the longer neural circuit - involving the auditory cortex - would trigger the motor response. This model would explain why responses are faster when SCM activity is observed. Interestingly, the StartReact effect can still be observed when the startle reflex is reduced due to the presentation of a less intense stimulus before the go-signal (pre-pulse inhibition, PPI) [15]. This result is counterintuitive because it suggests that the more direct neural circuit is still activated when the transient activation of the midbrain nuclei by the prepulse stimulus exerts long-lasting inhibition of the giant neurons of the caudal pontine reticular nucleus and inhibits the neural circuit of the startle reflex [42]. Alternatively, we have suggested that the apparent correlation between SCM activity and premotor RT could be a result of variations in the build-up of preparatory activity from trial to trial [16]. More precisely, it seems plausible that in some trials, the peak of preparatory activity could be either reached sooner than expected (and remain constant until the go-signal arrives) or at the expected time of the go-signal, whereas on other trials the go-signal could reach peak slightly later. If we assume that the build-up of preparatory activity during the foreperiod facilitates SCM muscle activity, then it could be the case that SCM+ trials simply indicate a higher level of preparatory activity, resulting in responses being initiated earlier when preparatory activity is relatively higher. This hypothesis would explain why responses with SCM activity are faster than those without it, eliminating the requirement for different neural circuits involved in response triggering. We experimentally tested the effect of varying the levels of preparation during the course of a trial on the latency of responses triggered by loud startling stimulus.

#### Experiment

To induce systematic fluctuations of the overall state of preparation for an action over the time course of a trial, we employed a marked reaction time task (see [29]). This task was expected to make participants alter the level of preparation for an action as a function of the conditional probability of the IS being presented in the unpredictable block of trials. More specifically, in any given trial, the probability that the IS would appear with the next time marker should increase, resulting in an increased level of preparation for action. In agreement with this prediction, RTs decreased linearly as the IS was presented later on a trial in the unpredictable blocks of trials (see Figure 6), and this was true for RTs to the soft and loud ISs (SAS). This effect was clearly reduced in the predictable block of trials, however, responses still seemed faster when the IS was delivered later in the trial (together with the last marker), and the probability of a false start increased linearly as the time between warning and IS time increased. Thus, our task successfully led participants to increase the overall level of preparation over time. To avoid the possibility that participants completely anticipated the time to initiate their action and respond to the IS only, we introduced catch trials in which no IS was presented and responses should be withheld. Our analysis of false starts showed that participants had more false starts in the predictable than in the unpredictable block of trials. This is an important observation because it indicates that when the IS is certain to appear and the foreperiod is predictable, participants are more likely to exhibit anticipatory reactions that can be mistaken for very fast responses to the IS in the context of the StartReact effect.

Interestingly, despite the fact that participants knew in advance the time of the IS in the predictable block of trials, the probability of falsely starting a response increased as the IS was presented later. This suggests that preparatory activity could still increase over time, and that participants do not simply engage in motor preparation after the 3rd IS marker (last 500 ms) in the predictable block of trials. If this is the case, RT is not really a sensitive measure of motor preparation when the foreperiod is predictable and the IS intense as RTs can be very close to the limit of the human capability. Note that the mean RT in the predictable block for the 3rd IS marker was 103 ms (% of false starts = 26), which would correspond to a premotor RT of about 84 ms (see our estimate of the neuromechanical delay in the Data Analysis section). This estimate would be 96 ms in the unpredictable blocks of trials, when the percentage of false starts was below 5%. To further examine whether preparation levels would affect the latency of responses to SAS, we calculated the 30th and 60th percentiles for these responses. The results of this analysis indicated that even the fastest responses to SAS are affected by rising preparation levels. Here the fastest RT in the predictable block of trials would correspond to a premotor RT of about 73 ms in the predictable block of trials (RT to the 3rd IS marker), and 86 ms in the unpredictable block of trials. These estimates are within the range of what one expects when analyzing the StartReact effect. Moreover, the difference between responses at the 30th and 60th percentiles, assuming a bimodal distribution of trials, was 21 ms in the predictable block and 18 ms in the unpredictable block of trials. These values are well within the expected range we obtained in our mini-meta-analysis (95% CI [-21.8, −9.3], see Figure 1) suggesting this might be a valid method to separate trials in the context of the StartReact effect where entire datasets are often discarded from analysis when participants do not display clear SCM activity.

The average slope of the linear regression function calculated using a bootstrapped procedure was −0.031 in the unpredictable block of trials, which means RTs could increase by up to 18.6 ms if one mistimed the peak of preparatory activity by 600 ms. This estimate is more than enough to account for a substantial proportion - if not all - of the RT advantage when SCM activity is detected via surface EMG. Of course, the slope was very close to zero, when the IS was highly predictable, but here responses were already so fast that it can be difficult to detect reliable effects so close to the neurophysiological limits of the system to react voluntarily to the IS. Thus, these results are in agreement with our hypothesis that the build-up in preparatory activity during the foreperiod does facilitate the RT of very fast reactions, and - as previously demonstrated by different groups [43-45] - this can increase the chances of observing SCM muscle activity.

## Conclusion

In conclusion, trials in which SCM activity is observed (SCM+) are faster than those without it (SCM-), with a medium to large effect size. Although the mean difference between SCM+ and SCM- trials is big enough to suggest separate neural pathways for movement triggering, the average premotor RT in SCM+ trials (simple RT tasks) is not short enough (≈90 ms) to rule-out cortical involvement. It is clear from our meta-analysis that foreperiod variability has a large impact on the average premotor RT on control trials (softer go-signal), making it impractical to determine whether the StartReact effect occurs based on the magnitude of the difference between SCM+ and control trials. Our experimental data demonstrated that responses initiated very quickly by SAS are still affected by the immediate level of preparation during a trial, and could partially - or completely - explain why SCM+ trials tend to be faster than SCM- trials. Our experimental task and approach to distinguish fast (StartReact) and slow (voluntary reactions) responses to SAS is a promising method to advance our knowledge about the mechanisms involved in the preparation and initiation of motor actions.

## Acknowledgements

This work was funded by Australian Research Council Discovery Project Grant DP160102001.

